# Vocal distinctiveness in young Eurasian Scops Owls (*Otus scops*)

**DOI:** 10.1101/2023.09.17.558098

**Authors:** Fabrizio Grieco

## Abstract

Vocal distinctiveness is expected to occur more often in colonial-breeding species as the parents need to recognize their offspring in a large group of conspecifics. Territorial species like the Scops Owl are expected to exhibit low distinctiveness. Contrary to what was expected, spectrographic analysis of the food-begging calls of young Scops Owls revealed previously unknown, complex acoustic structure.

Within recording sessions, call duration exhibited the highest repeatability (average R 0.82), followed by the peak frequency in the last third of the call (0.61). Other spectral measures showed low to moderate repeatability (0.32 – 0.57), while the time between subsequent calls was the least repeatable (0.15). When comparing recordings made on different nights, Linear Discriminant Analysis assigned 55.7% of the calls to the correct individual, and 73.1% when restricting analysis within broods. When analyzing variability across recordings, individuals explained most of the variation in Call duration and Peak frequency in the last third of the call (89.5 % and 81.2 %, respectively), while recordings explained little variation (3.4 % and 1.3 %, respectively), suggesting that those acoustic features were the most important in vocal stability and distinctiveness. The calculated information capacity H_S_ was 4.48 bits, i.e. within the range of values found in loosely-colonial species. The results suggest that the vocalizations of young Scops Owls show moderate individuality that could only help offspring recognition among a small number of individuals.

However, the functional significance of call distinctiveness remains unclear; a few hypotheses are discussed. *Keywords*: Acoustic signature, coloniality, fledglings, individuality, Otus scops, repeatability.

## INTRODUCTION

The ability to discriminate among conspecifics is an important characteristic in many animals and is often required for mate and kin recognition, and parent-offspring communication (Tibbetts & Dale 2007).

Studies have cumulated evidence that vocal distinctiveness is widespread across taxa including birds (Falls 1982, Lambrechts & Dhondt 1995, Carlson et al. 2020), and show the importance of distinctiveness in contexts such as mate and kin recognition (e.g. Nakagawa & Waas, 2004; Sharp et al., 2005; Vignal et al., 2008) neighbour-stranger discrimination (Stoddard 1996), and parent-offspring communication (Beer 1971, Tibbetts & Dale 2007). Mechanism of recognition between parents and their offspring has often been found in bird species that breed in dense and noisy colonies, like penguins and seabirds (Buckley & Buckley 1972, Lefevre et al. 1998, Jouventin & Aubin 2002). In colonial swallow species, nestlings make individually distinctive calls that parents recognize (Beecher et al. 1981, Stoddard & Beecher 1983, Loesche et al. 1991, Strickler 2013). On the contrary, calls of nestlings of non-colonial species show weak individuality (Medvin & Beecher, 1986; Medvin et al., 1992, 1993; Leonard et al. 1997; but see Barg & Mumme, 1994). The observation that parent-offspring recognition is better developed in colonial than in noncolonial species led to the hypothesis that coloniality is a strong selection pressure for the evolution and maintenance of the distinctiveness of calls (Beecher 1990). In colonially breeding species, the parents must identify their own young among many unrelated ones, particularly after their young leave the nest and become mobile, so that their location is an unreliable indicator of identity. In those cases, the risk of misdirected parental care is high.

Individual variation in vocalizations is a prerequisite for the ability to discriminate between conspecifics provided that visual or other types of cues are not available (Falls 1982), and when the parents need to find their offspring in a crowded area far from the nest. However, vocal traits often vary along with the physical or behavioural conditions of the individual, so to be a reliable indicator of identity, a signal must also be temporally stable, that is, it must remain unaltered along significant time scales (Gilbert & McGregor 1994, Peake et al. 1998, Lengagne 2001, Grava et al. 2008, Klenova et al. 2009, 2012). Among *Strigiformes*, vocal signatures have often been found in adult birds (e.g. Eagle owls: Lengagne, 2001; Great horned owls, Odom et al., 2013; Barred owls, Freeman, 2000; Tawny owls, Galeotti & Pavan, 1991; Pigmy owl: Galeotti et al., 1993; Western screech-owls, Tripp & Otter, 2006; Ural owl, Zhou et al., 2020; Northern spotted owl, Fitton, 1991; Eastern screech owl Cavanagh & Ritchison, 1987), suggesting that identity signals are favoured in this group of birds, particularly in the context of territorial interactions and communication with the breeding partner. In contrast with the ample evidence of individuality of adult vocalizations, little is known about individuality in earlier life stages, with some well-known exceptions, for example in penguins, seabirds, and swallows (Carlson et al. 2020). In owls, the only species where individual variation of juvenile calls has been established is, to date, the Barn Owl (Dreiss et al. 2014).

The Eurasian Scops Owl (*Otus scops*, hereafter ‘Scops Owl’) is a small (body length 19-20 cm), nocturnal, primarily insectivorous owl, that breeds in open habitats in the Mediterranean zone and Central-Eastern Europe (Mikkola 1983, Cramp 1985). The species is mainly territorial, although it sometimes forms “loose colonies” of breeding pairs with inter-nest distances as short as 15 m (Grieco 2018; for an overview see Cramp 1985). The young (in the study area, one to four per brood) leave their nest about 21 days after hatching (Koenig 1973), and on the following nights move further and further away from the nest. The parents keep feeding the young regularly up to 40 days after hatching, then increasingly less often. At 45 days the young are fully able to catch prey (Koenig 1973), and become gradually independent and disperse.

Since the first day of their life, young Scops owls produce short (0.15 s) “tschp” calls at intervals of 0.5-1 second, interpreted as food solicitation calls (Koenig 1973). Besides the spectrograms reported by Koenig (1973), there is no quantitative data available on the call structure, and very little is known about the development of the vocalizations, their inter-individual variation, and their stability over time. Koenig (1973) suggests that the begging call gradually transitions during development to the advertising call of the adults. Because the call of the adult owls is individually distinctive (see below), looking at the calls of the juveniles is also important in the context of the development of individuality.

When formulating hypotheses around the function of vocal distinctiveness early in life, between-species variation in individuality is often linked to breeding biology. The Scops Owl is predominantly territorial, meaning that, after fledging, the young do not join large groups of conspecifics that pose the problem of parent-offspring recognition. This leads to the expectation that the parents do not need to recognize their offspring individually after they leave the nest. Therefore, the degree of individuality in the vocalizations of the young should be low, within the range found in other species that are non-colonial or breed in small groups (Medvin et al., 1993).

This observational study had two aims: (1) To provide the first detailed description of the food-begging calls of young Scops Owls and their inter-individual variation based on spectrographic analysis; (2) To determine whether individual owlets produce calls that are consistent in their structure and temporally stable. Information about vocal distinctiveness in the post-fledging phase is particularly relevant since most of the studies of vocalizations and begging behaviour of young birds have focused on nestlings, whereas the post-fledging phase is underrepresented (Draganoiu et al. 2006, Middleton et al. 2007), particularly for nocturnal bird species (Kouba et al. 2014).

## METHODS

### Study area

The study was carried out at the State Reserve Metaponto, Italy (40°22’24“N, 16°50’19’’E), a reforested area dominated by Aleppo pine *Pinus halepensis* along the sea coast. Forest plots are mixed with human settlements, especially recreational resorts and camping parks. The Scops Owl is a primarily cavity-nesting species; however, tree cavities are virtually absent in this area. Breeding pairs occupy old twig nests of Magpies (*Pica pica;* Grieco 2018).

### Sound recording and call selection

I recorded vocalizations of 20 Scops Owl nestlings and fledged young from six broods between July and August 2021 and July 2022. Recording sessions were carried out between 19:30 and 21:30 and between 00:00 to 04:30 and were made opportunistically. The study area was actively searched for Magpie (roofed) nests and fledged broods. Previous experience indicated that Scops Owls avoid twig nests completely open as breeding sites. Magpie nests were watched in the evening and in the morning for at least half an hour to detect adult owls bringing food. Fledged young were found by listening to their calls at a distance. To record sound, I used a Telinga Pro 8 MK2 stereo microphone mounted on a parabolic dish with a diameter of 52 cm, and a Marantz PMD 661 MK2 digital sound recorder. By pointing the microphone toward a vocalizing subject, it was possible to amplify its calls while keeping the voice of its siblings in the background. This allowed to track the owls unobtrusively but at close range. The beeline distance between the microphone and the owls ranged between four and 15 meters. In addition, because siblings were roughly at the same distance to the microphone, the large differences observed in the spectral structure were unlikely to be due to distance-related attenuation of higher frequencies.

Furthermore, there was no evidence that the recording procedure altered the birds’ behaviour: (1) The adults gave alarm calls only when they detected cats or foxes around the nests. As soon as they did so, the young stopped calling. The fact that I could record so many calls at short distances indicates that the adults were not stressed. (2) The fledglings stayed in the same tree or area for several minutes to hours, and the adults kept bringing food to the nest or fed the young regularly during the recording sessions.

Each recording lasted at least one minute and was stored in a WAV file at a sampling rate of 44.1 kHz, and with 16-bit depth. In most cases, recordings were made a few meters from the nest tree or the fledged young.

Recordings lasted on average 1235 ± 1018 (SD) seconds (range 80-3900). The total length of the recordings was 16 hours 28 minutes. Because hatching was not observed directly, the exact age of the nestlings was unknown. Following Koenig (1973) fledging was estimated to occur on average 24 days after hatching (range 21-30). For four of the six broods, the fledging date was known, and the first recording was made two days before fledging to 10 days after fledging (see Supplementary Table S1). For the other two broods with an unknown fledging date, their frequent movements observed during recording sessions and the low visit rate by the parents suggest that the young might have left their nest one to two weeks earlier.

For each of the 23 individuals and each recording, I selected the 10 loudest calls that were free of background noise and did not overlap calls of their mates or adults. In total, 990 calls were analysed. Calls were assumed to be of the type “food-begging *sensu* Koenig (1973). Other utterances like bill-snapping and trills that mostly occurred during parental visits were ignored. Calls were selected that occurred at least 30 s before and 30 s after a parental visit. That criterion was adopted because previous observations suggested that young owls altered their calls’ structure and rhythm in the presence of the adults (see Results). An effort was made to select calls that covered the entire recording length. However, this was not always possible for two reasons: (1) the young sometimes produced calls too faint to be reliably measured, and (2) part of the recording contained higher levels of background noise, primarily due to the choruses of cicadas and other insects and human-induced noise. All sorts of wide-band noise made measurements of some acoustic features like Center of gravity and Entropy unreliable. Therefore, the calls selected were not equidistant. Overall, the median time between consecutive calls within a recording was 19.2 s (interquartile range 4.4 - 62.5 s).

The calls were assigned to individuals after visually comparing the spectrograms with those obtained from previous recordings of the same brood. Individual spectrogram patterns occurred in “discrete” forms, in most cases easily recognizable. When the owlets were detected visually (seven out of eight broods), there were always as many forms as the number of owlets observed. However, assigning IDs based on call similarity and later claiming that the calls were repeatable within individuals is open to circular reasoning. When juveniles were acoustically separated (single fledglings, or fledglings that were temporarily far from their siblings) they produced calls of one consistent form and relatively stable acoustic structure throughout the recording time, even over multiple days (see Results, Figure 1, and Supplementary Information S1). Furthermore, if individuals had switched to other spectrogram forms from time to time, there should have been a high chance of observing two individuals making calls of similar structure. However, that was never observed. This suggests that the chance of mistakenly mixing the “true” identity of the brood mates was reasonably low.

**Fig. 1.**
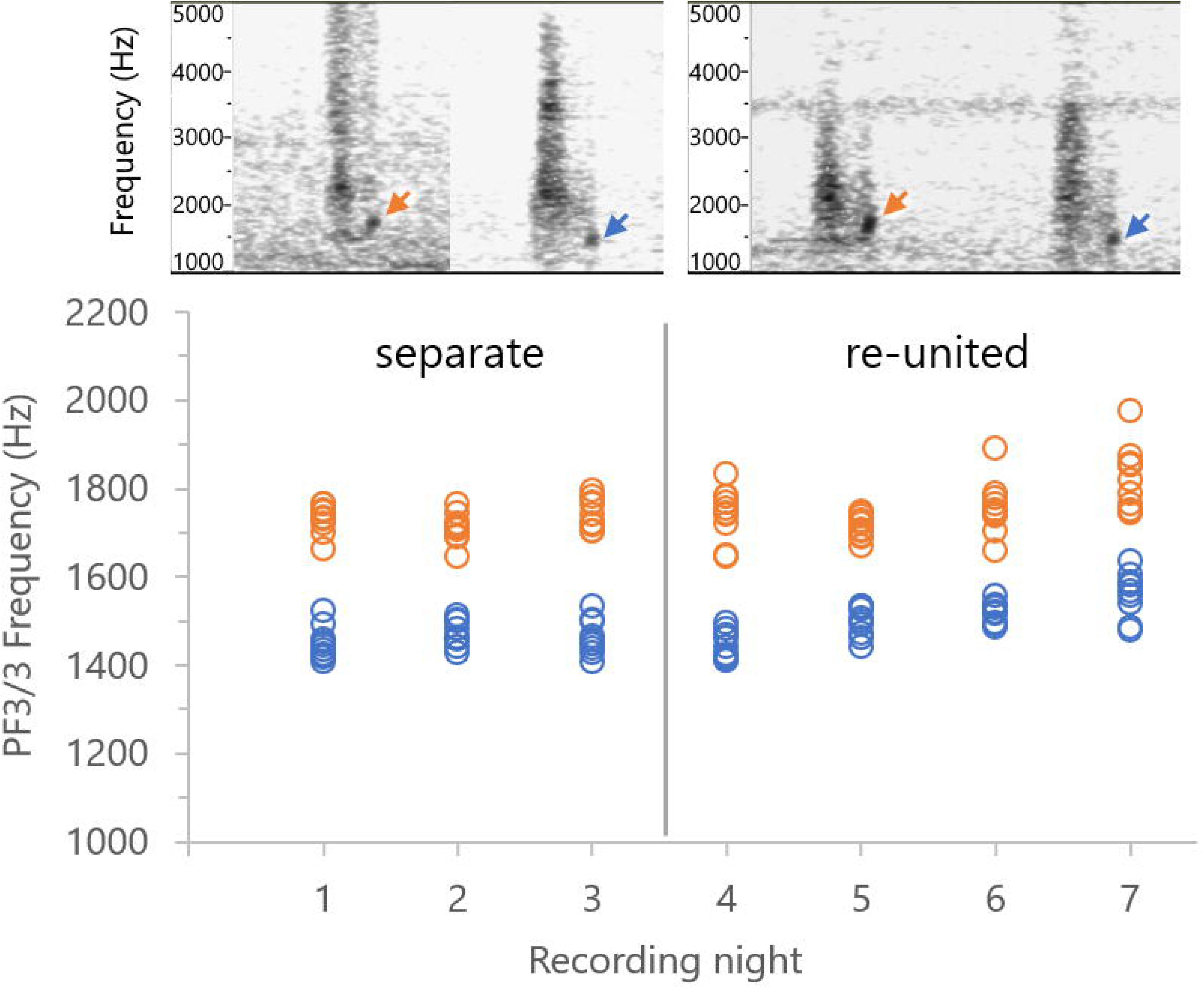
Evidence that acoustic features of calls can be used to track individual Scops Owl fledglings. Representative spectrograms of the calls of two siblings when they perched at a distance for three subsequent nights (left) and then reunited (right). The arrows indicate the peak frequency at the end of the call (PF3/3, see Methods) used to separate the individuals. The total time step is one second. Bottom: ten measurements of PF3/3 for each individual, on each night. The colors indicate the individual classified based on the visual inspection of the spectrogram and the perceived sound.

To analyse the effect of parental visits on the call’s acoustic properties, I selected 10 random calls occurring at least 30 s before or 30 s after a visit (“Random calls”) and I compared them with the calls that occurred less than 5 s before a visit, or up to 10 s after a visit (“Visit calls”). Visit calls of siblings were often overlapping, so it was possible to select vocalizations only for a small number of individuals. This resulted in a smaller sample of individuals for this analysis (*n* = 12).

### Acoustic analysis

Sound analysis was carried out in Audacity 2.1 (https://www.audacityteam.org), Praat 6.0 (https://www.fon.hum.uva.nl/praat/), and Sound Analysis Pro (http://soundanalysispro.com/) depending on the acoustic features examined. Recordings were first filtered with a 1000-Hz high-pass filter in Praat 6.0 to remove low-frequency background noise. The following FFT settings were used: FFT window size 2048 samples, window type Hanning, temporal resolution 46 ms, frequency resolution 21.5 Hz. Two temporal features and seven spectral features were measured (for details see Call features in Supplementary Information 1):

1. Call duration (s). The time from the onset to the end of the call, measured from the waveform.
2. Time from the previous call (s). The interval between the onset time of the previous call and that of the current call, measured from the waveform.
3. Peak frequency 1/3 (PF1/3, Hz). The frequency at maximum amplitude within the first third of the call duration.
4. Peak frequency 3/3 (PF3/3, Hz). The frequency at maximum amplitude within the last third of the call duration.
5. Centre of gravity (Hz). The average frequency over the entire frequency domain, weighted by the sound amplitude.
6. Pitch (Hz). The average pitch across the call duration. The intended use of Pitch was to describe the perceived tone of a sound.
7. Average entropy. The average Wiener entropy across the call duration. Entropy measures the extent to which the sound contains a mixture of frequencies, as opposed to a single frequency or tone.
8. Minimum entropy. The minimum Wiener entropy within the call duration. It is usually found at the end of the call.
9. Spectrum skewness. A measure of the symmetry of the spectrum. If skewness is zero, the spectrum is perfectly symmetrical.

### Statistical analysis

*Repeatability*. To assess the repeatability of acoustic features within broods, I selected the recordings of the broods with at least two brood mates (n = 7). For each acoustic feature, I performed a one-factor ANOVA for each recording with young ID as a factor and calculated the Repeatability (*R*) using the formula by Lessells & Boag (1987). The data set included 630 calls. Broods were represented with a variable number of recordings, one to seven. Each combination brood*recording resulted in one ANOVA and thus one value of R, for a total of 21 R estimates for each acoustic feature. The average R was calculated from the per-brood averages. 95% Confidence intervals were calculated from following Nakagawa & Schielzeth (2010).

Data sets subject to ANOVAs were tested for normality. For each recording, I considered the residuals of the measurements after fitting the individual means (i.e., each measurement diminished by the mean within the recording). Kolmogorov-Smirnov tests were applied to the sets of 10 measurements (48 sets for each of the nine acoustic features). After Benjamini-Hochberg correction for multiple testing (FDR 0.05, n = 48 tests), the null hypothesis of normality was rejected in six out of 48 sets for Peak Frequency 1/3 and in one of 48 sets for Peak Frequency 3/3.

*Proportion of variance*. To assess the amount of variance accounted for by individual birds and recordings, I analysed the broods that were measured at least two times (broods 4 to 8, n = 13 young). To account for multiple measurements of the same individual and the different number of recordings available, I applied linear mixed models in R (R Core Team 2023) using the *lme4* package (Bates et al. 2015) where individual and recording were random factors. P-values were obtained from Likelihood Ratio tests of the model with the effect in question against the model without that effect. The proportion of variance explained by a factor was calculated from the figures given by the summary() function of the full model. The model was run for each acoustic feature. Brood was considered a fixed factor because of the small number of levels (i.e., 3), and was removed from the final model for all acoustic features as it was not significant. Data was checked for normality using Q-Q plots of residuals and residuals were plotted against the variables under study. Furthermore, the estimates of the random intercepts (eblups) were visualized in Q-Q plots and histograms. Variables with skewed distributions were transformed using inverse or log transformation (inverse 1/x: Centre of gravity, Peak frequency 1/3 and Peak frequency 3/3; log_10_(x): Call duration and Time from the previous call).

*Linear Discriminant Analysis*. Another way to look at the stability of individual differences in vocalizations is to use Linear Discriminant Analysis (LDA). This tests whether a specific call can be attributed to an individual based on the acoustic features of calls recorded in a different (often earlier) session. To keep a balanced design with an equal number of calls per individual, I selected 10 calls per recording, per individual, for the broods that were recorded at least two times. Of the nine acoustic features, seven were entered as explanatory variables. The other two, Time from the previous call and Average entropy, were excluded because the former showed low repeatability (see Results) while the latter was highly correlated with Minimum entropy (see Table 2). Acoustic features had different variances due to the different scales. Because LDA is sensitive to the relative size of the features, each feature value was standardized by removing the average and dividing by the standard deviation. LDA was performed in XLSTAT for Excel (Lumivero 2023). In the first LDA, two recordings separated by 1 to 3 days were selected to obtain a homogeneous dataset. The earlier recording was used to compute the discriminant function while the later recording was used for validation. Each individual (n = 14) was represented by twenty calls; 10 for training the discriminant function and 10 for validation. In the second LDA, the stability of the calls was tested for a longer period (6-9 days difference between training and validation; n = 5). In both cases, LDA was run twice: across all broods (i.e., each call in the validation set was tested against the calls of all individuals in the training set) and within each brood (i.e., each call in the validation set was tested among the calls of the brood mates in the training set). Classification success (CS) was the percentage of calls classified correctly.

One important limitation of the above-mentioned CS is that it much depends on sampling; its meaning is limited to the specific number of individuals used. In general, CS decreases with larger samples (Linhart et al. 2019) making comparisons between studies difficult. Beecher (1989) proposed the use of a statistic derived from the theory of information, H_S_, the estimated number of bits that can describe an identity signal. From H_S_, one can directly derive the size 2^Hs^ of the group in which individuals can be identified with a certain degree of accuracy (Beecher 1989). Although H_S_ is not completely independent of sampling (Linhart & Šálek 2017), it can be considered a population estimate and enables meaningful comparisons between studies and across species (Linhart et al. 2019). To calculate H_S_ following Beecher (1989), I used the variables that exhibited significant inter-individual variation after standardization (i.e., each variable was divided by its standard deviation estimate σ_w_) and Principal Component Analysis transformation. After performing an ANOVA on each PCA factor (effect of individual bird, all P < 0.0001), the individual values H_S_ obtained from the between and within mean squares were summed up to give the statistic H_S_.

## RESULTS

### Evidence that different spectrogram patterns represent different individuals

Because young owls were not marked, a way to prove that different spectrograms reflected the voices of different individuals came from analysis of the calls of siblings that were temporarily spaced apart. Two fledglings of one brood were found at different locations during three nights; one of them was found near the natal nest while the other, having left the nest earlier, perched at about 20 m distance. The two owlets produced calls consistently different at the visual inspection of the spectrogram and in an acoustic feature, Peak frequency 3/3, associated with the “dot” pattern in the terminal segment of the calls (Figure 1, top). One could object that the two owlets may have exchanged their position between subsequent recordings and each produced the same vocalization that its brood mate made in the previous recordings. However, that seems an unlikely scenario. Eventually, the two owlets re-united and stayed together for a few days, until one disappeared. The calls recorded after the reunion showed the same difference in PF3/3 during multiple nights (Figure 1). Similarly, two fledglings of another brood were recorded at about 5-10 m distance from each other for half an hour. Because the microphone was pointed to each individual at specific times, one could tell from the relative amplitude of the calls which animal produced a specific call. The two individuals showed consistent differences in call duration. When the calls were labeled based on the spectrogram patterns only, the differences in duration were still present across multiple nights (Suppl. Figure 1, A vs. B). This demonstrates that different spectrogram forms, exhibiting consistent differences in acoustic properties, indicated the presence of different individuals. Other examples are reported in Supplementary Information S1.

### Basic statistics and covariation between acoustic features of the calls

Calls of young Scops Owls showed marked variability in all the temporal and spectral features measured (Table 1). The first part of the call is generally noisier, with a wide spectrum and peak frequency of up to 4000 Hz in most cases (Peak frequency 1/3). The last part is more structured, with a narrower, more asymmetric spectrum, with the loudest frequency components being under 2000 Hz (Peak frequency 3/3). This is the fraction of the call where entropy is the lowest. Visual inspection revealed a high degree of variability of the call spectrogram within broods, in the form of dot-and S-shape patterns (Figure 1, Figure 2). In Supplementary Video S3 one can compare the calls of four brood mates and notice the differences in both the spectrogram form and the corresponding perceived sound.

**Fig. 2.**
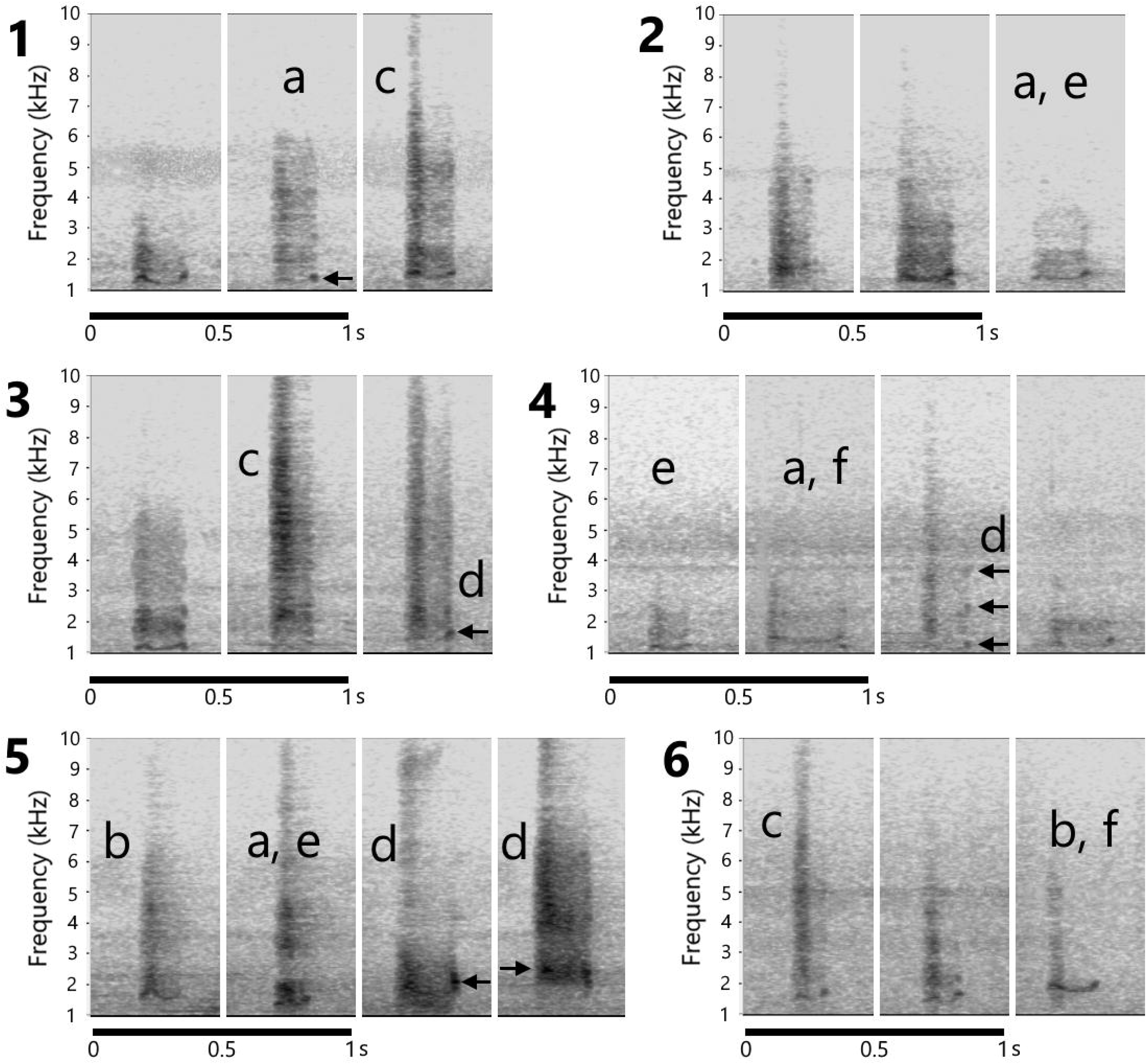
Representative spectrograms of the calls of young Scops Owls of six broods. Letters indicate repeatable features based on visual inspection. (a) Differences in call duration. (b) S-shape patterns at the lower end of the frequency range. (c) Wider frequency band in the first part of the call, resulting in a higher Centre of gravity and lower Spectrum skewness. (d) Acoustic energy concentrated at specific frequencies (indicated with arrows). (e) The call sounds lower in pitch than those of brood mates. (f) More structured sound, with high spectral skewness.

**Table 1.**
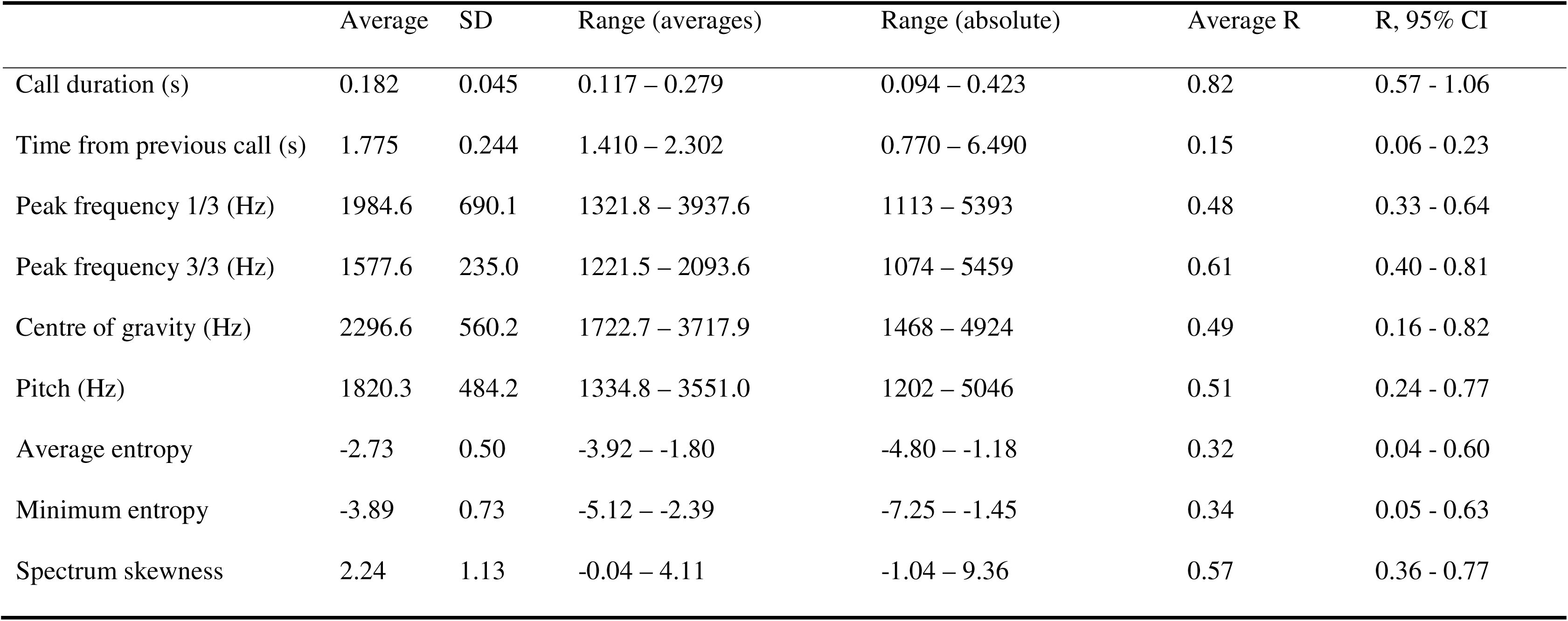
Descriptive statistics of acoustic variables of vocalizations of young Scops Owls. The Range is calculated using individual averages (n= 23) and using single measurements (n= 990 calls). Estimates of repeatability R (average and 95% confidence intervals) are based on broods of two or more fledglings (n = 7 broods, 22 individuals, 630 calls).

**Table 2.** Pearson’s correlation coefficients of temporal and acoustic variables in calls of 20 young Scops Owls, based on the per-individual averages. * P < 0.05; ** P < 0.001. Values in bold indicate a significant correlation after Benjamini-Hochberg correction (FDR = 0.05, n = 36 tests).

|  | Time from<br>prev. call | Peak freq. 1/3 | Peak freq. 3/3 | Centre of<br>gravity | Pitch | Average<br>entropy | Minimum<br>entropy | Spectrum<br>skewness |
| --- | --- | --- | --- | --- | --- | --- | --- | --- |
| Call Duration | 0.32<sup>ns</sup> | -0.44<sup>ns</sup> | 0.07<sup>ns</sup> | **-0.53 \*** | -0.30<sup>ns</sup> | -0.09<sup>ns</sup> | -0.23<sup>ns</sup> | **0.50 \*** |
| Time from prev. call |  | -0.10<sup>ns</sup> | 0.39<sup>ns</sup> | -0.11<sup>ns</sup> | -0.11<sup>ns</sup> | -0.01<sup>ns</sup> | -0.11<sup>ns</sup> | 0.36<sup>ns</sup> |
| Peak frequency 1/3 |  |  | 0.47 * | **0.91 \*\*** | **0.81 \*\*** | 0.31<sup>ns</sup> | 0.35<sup>ns</sup> | **-0.77 \*\*** |
| Peak frequency 3/3 |  |  |  | 0.46 * | **0.54 \*** | -0.06<sup>ns</sup> | -0.16<sup>ns</sup> | -0.20<sup>ns</sup> |
| Centre of gravity |  |  |  |  | **0.81 \*\*** | 0.43 * | 0.46 * | **-0.90 \*\*** |
| Pitch |  |  |  |  |  | 0.16<sup>ns</sup> | 0.19<sup>ns</sup> | **-0.62 \*** |
| Average entropy |  |  |  |  |  |  | **0.92 \*\*** | -0.41 * |
| Minimum entropy |  |  |  |  |  |  |  | **-0.50 \*** |

**Table 3.** Results of Linear Mixed Models of acoustic features of 860 calls of young Scops owls in five broods: brood 4 (four young, two recordings over four days), 5 (four young, five recordings over five days), 6 (two young, 13 recordings over 10 days), 7 (two young, seven recordings over seven days) and 8 (one young, nine recordings over nine days). Effects were tested with Likelihood Ratio after comparison between the model with both random factors and the model with the other factor only. The effect of brood was not significant for any features (always P > 0.05). (*) Significant after Benjamini-Hochberg correction for multiple testing (FDR = 0.05, 9 tests).

|  | Effect: Individual |  | Effect: Recording |  | Residual variance (%) |
| --- | --- | --- | --- | --- | --- |
| | $\chi^2$, df = 1 | Prop. of variance (%) | $\chi^2$, df = 1 | Prop. of variance (%) | |
| Call duration | 1648.7 * | 89.5 | 239.7 * | 3.37 | 7.11 |
| Time from prev. call | 49.9 * | 11.49 | 76.1 * | 21.15 | 67.36 |
| Peak frequency 1/3 | 169.2 * | 43.07 | 37.1 * | 5.27 | 51.66 |
| Peak frequency 3/3 | 558.7 * | 81.22 | 54.6 * | 1.32 | 17.46 |
| Centre of gravity | 316.4 * | 35.87 | 140.4 * | 17.12 | 47.01 |
| Pitch | 215.4 * | 43.42 | 99.7 * | 11.67 | 44.91 |
| Average entropy | 141.6 * | 21.58 | 193.6 * | 29.29 | 49.14 |
| Minimum entropy | 303.4 * | 29.58 | 244.1 * | 27.87 | 42.55 |
| Spectrum skewness | 319.0 * | 29.97 | 139.7 * | 16.18 | 53.86 |

Several correlations were found between the calls’ acoustic features (Table 2). Frequency measurements co-varied in general with each other, as expected. However, Peak frequency 3/3 did not correlate with any other measure, except for a weak correlation with Pitch. Young owls that produced higher-frequency calls also produced calls with a more symmetric and broad spectrum, as indicated by the negative relationship between Peak frequency 1/3, Center of gravity, Pitch, and Spectrum skewness (Table 2). Longer calls had a lower center of gravity and were more skewed. However, the time from the previous call, which reflected the calling rhythm of the individual, was not associated with any other measure.

### Distinctiveness of calls

Repeatability estimates within broods were significantly greater than zero for all the features measured (Table 1). Call duration showed the highest repeatability, suggesting that it was the best indicator of individual distinctiveness within broods. Frequency features and Spectral skewness showed moderate repeatability, the highest of which was that of Peak frequency 3/3. Average and Minimum entropy were less repeatable, while the time from the previous call exhibited the lowest average repeatability.

When addressing variability across multiple recordings, linear mixed models revealed that all call features exhibited significant effects of recording and individual (Table 4). Call duration exhibited the highest degree of variation due to subject identity (89%). For example, brood mates differed in their calls’ duration across multiple days (Figure 4). In contrast, Time from the previous call exhibited a high proportion of unexplained variance (67 %), indicating that the calling rhythm also varied greatly within recordings.

Measures of acoustic frequency gave more ambiguous results, with subject identity explaining a proportion of variance above 35 % (Table 4). However, Peak frequency 3/3 stood out with a high proportion of variance due to individuals, a lower proportion of variance due to recordings, and a lower residual variance. Figure 4 shows an example of the temporal stability of Peak frequency 3/3. Conversely, Peak frequency 1/3 which measured the sound frequency in the first part of the call exhibited a much lower proportion of variance due to individuals and a much higher proportion of residual variance. This suggests that the last part of the call contributed to individuality, while the first part of the call was too variable to reliably carry individual information.

Last, Spectral entropy and skewness showed a low to moderate proportion of variance explained by individuals and recordings (16 % to 30 %) and a greater amount of residual variance. This indicated that the degree of “noisiness” of the calls also varied between and within recordings.

Linear Discriminant Analysis aimed at classifying individuals (n = 14) based on the call features extracted from a previous recording session. On average, 55.7 % of the calls were correctly assigned to an individual based on the calls recorded one to three days earlier. The CS score was higher than would be expected by chance based on 14 individuals (Table 2 in Supplementary Information S1). When LDA was repeated within broods, CS increased to 73.1 %. This was partly expected since this time the calls had to be assigned to individuals within smaller groups of size 2, 3, or 4, depending on the brood an individual belonged to, instead of 14. However, CS was still higher than that expected from chance alone (Table 2 in Supplementary Information S1).

Subsequently, LDA was performed on a sub-sample of young that were followed for a longer time, i.e. the difference between the first and last recording was 6 to 9 days (n = 5). There, CS was 62.0 % when LDA was done across broods and 80.0% when repeated within broods. Again, CS was higher than expected by chance alone.

Analysis of the call features of 23 individuals indicated an information capacity H_S_ of 4.48. The largest contribution to this value (F1, H_s_ = 1.27), which expressed Centre of gravity (correlation between axis and variable: *r* = 0.95), Peak frequency 1/3 (*r* = 0.81), and Spectrum skewness (*r* =-0.84). This factor reflected the position of the highest amplitude along the frequency range, especially at the onset of the call. The second largest contribution (F2, H_s_ = 0.99) expressed Minimum entropy (*r* = 0.78), Peak frequency 3/3 (*r* =-0.76) and Average entropy (*r* = 0.69). Entropy usually dropped to the minimum at the end of the call, while PF3/3 described the peak frequency in the same part. Therefore, F2 reflected the amount of acoustic structure at the end of the call.

### Effect of parental visits on call structure

Young owls in the post-fledging phase vocalized continually, also in the absence of the parents. It was therefore possible to compare the acoustic structure of the calls in the absence of the parents with that during parental visits. Figure 3 shows the changes in the calls’ acoustic structure between random calls in the absence of the parents and those recorded around a visit. Compared with random calls, calls around parental visits were (1) shorter, (2) more frequent as the time from the previous call diminished, (3) generally higher in acoustic frequency, with a higher Centre of gravity and a lower Spectrum skewness, and (4) noisier, i.e., with higher entropy (Figure 3). However, Peak frequency 3/3, a relatively more stable feature that contributed to individuality, and Pitch did not significantly change during parental visits.

**Fig. 3.**
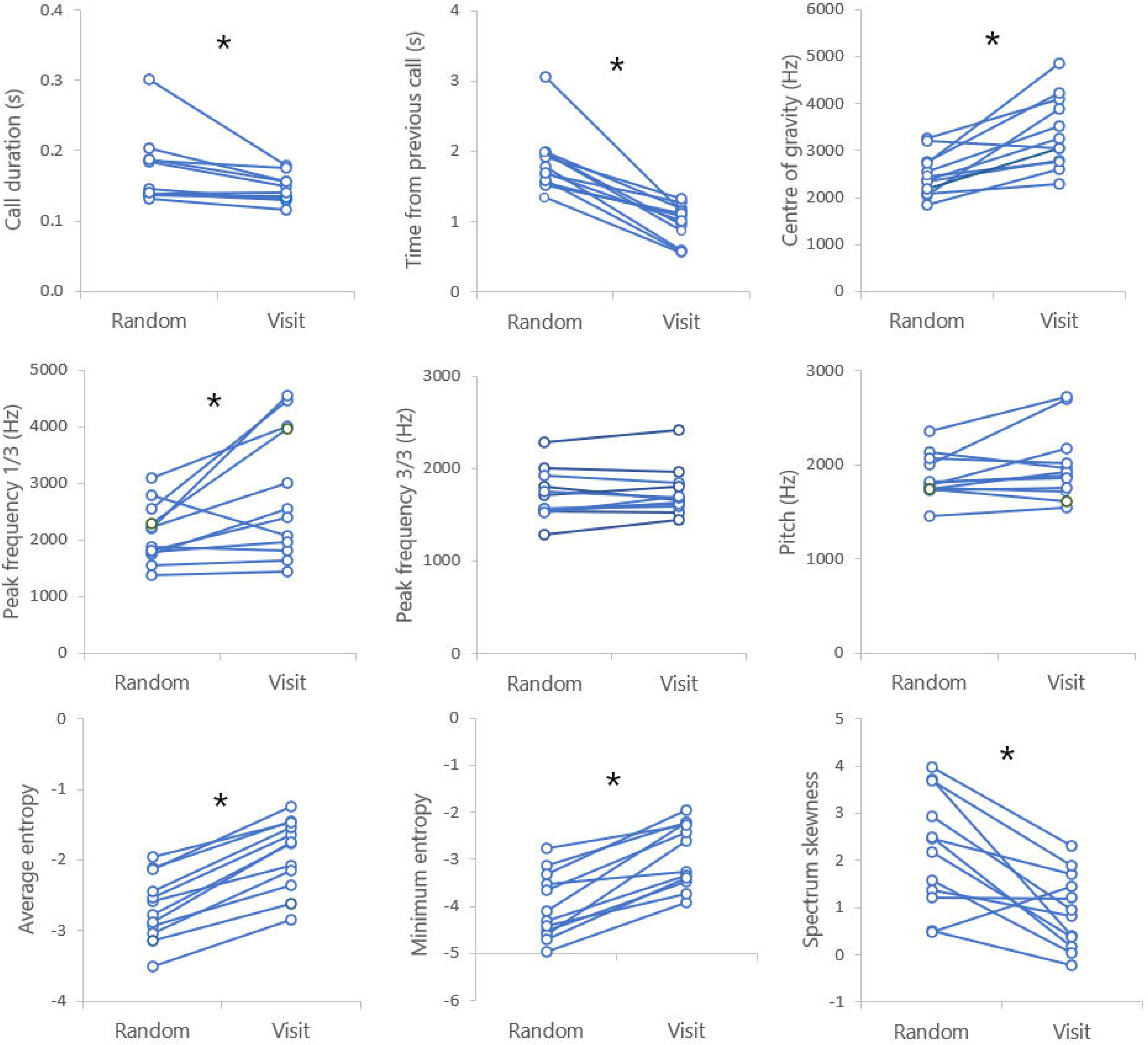
Within-subject changes in acoustic variables of the calls of young Scops Owls (all after fledging) during a parental visit. Each pair of circles represents one subject. Random: The average value over ten random calls at least 30 seconds before the visit and 30 seconds after a previous visit. Visit: The average of calls between 5 seconds before and 10 seconds after the visit. Asterisks indicate statistical significance after Benjamini-Hochberg correction of multiple testing (False Discovery Rate set to 0.05, 9 paired t-tests). Significant differences based on paired t-tests (Random vs. Visit, average ± SD, *n* = 12 individuals): Call duration, 0.17 ± 0.05 vs. 0.14 ± 0.02, *t*_11_ = 2.57, Cohen’s *d* = 0.74; Time from the previous call, 1.84 ± 0.44 vs 0.98 ± 0.27 s, *t*_11_ = 7.00, Cohen’s *d* = 2.02; Centre of gravity, 2487.4 ± 441.1 vs. 3363.0 ± 764.9 Hz, *t*_11_ =-4.56, Cohen’s *d* = 1.32; PF1/3, 2106.0 ± 512.9 vs. 2818.6 ± 1138.7 Hz, *t*_11_ =-2.76, Cohen’s *d* = 0.80; Average entropy,-2.70 ± 0.47 vs-1.91 ± 0.50, *t*_11_ =-12.29, Cohen’s *d* = 3.55; Minimum entropy, - 4.00 ± 0.70 vs-2.90 ± 0.68, *t*_11_ =-7.52, Cohen’s *d* = 2.17; Spectrum skewness, 2.20 ± 1.21 vs 0.93 ± 0.80, *t*_11_ = 3.91, Cohen’s *d* = 1.13; PF1/3 and Pitch n.s.

**Fig. 4.**
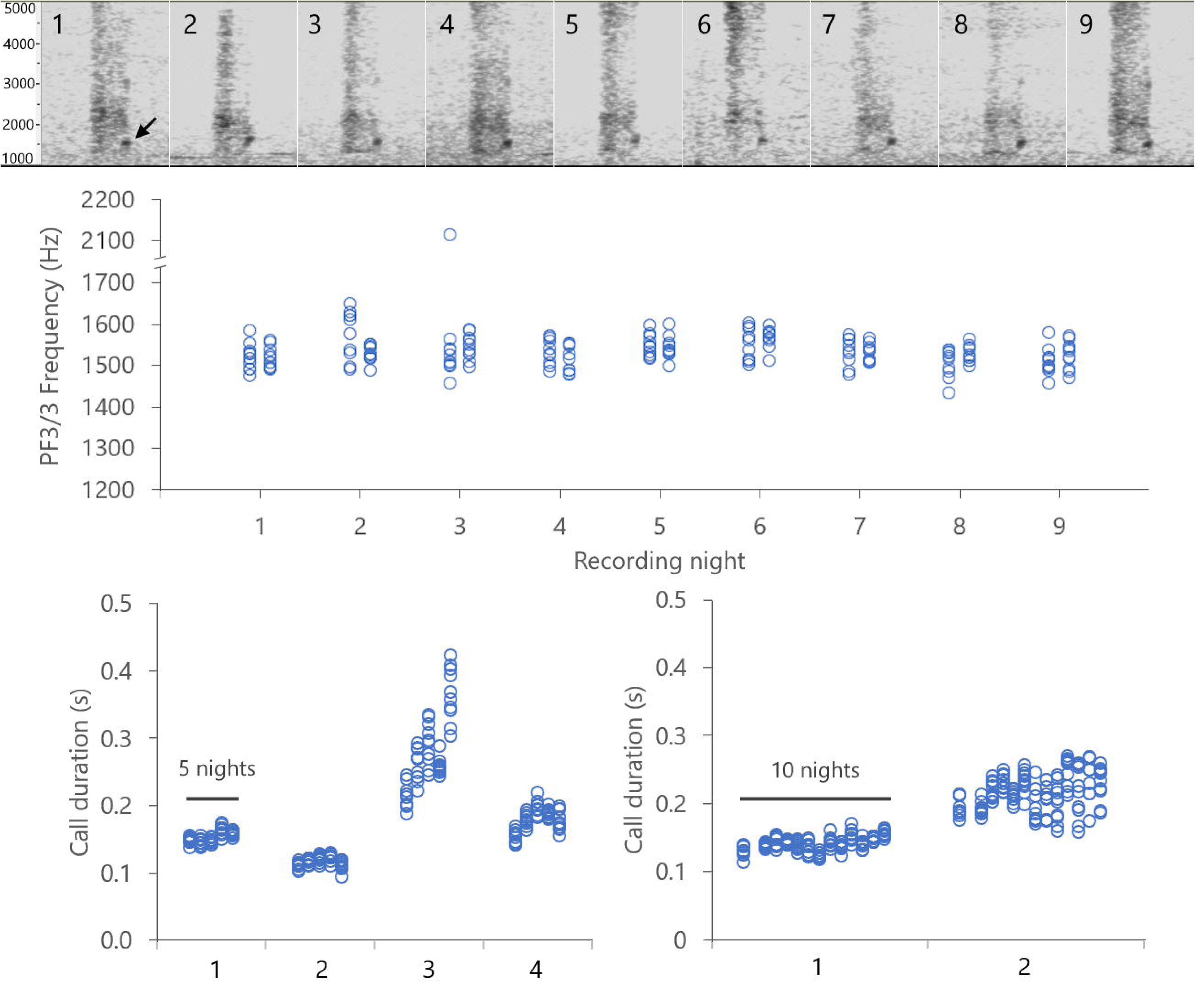
Temporal stability of acoustic features of calls of young scops owls. Top: representative spectrograms of the calls of a single fledgling (i.e., with no brood mates) in nine consecutive nights. Middle: PF3/3 of the calls of the same fledgling (two separate recordings per night, 10 calls selected per recording). Bottom: Call duration of fledglings in two broods, one with four individuals (left) and one with two (right). The horizontal bars indicate the number of days covered by the measurements. Each set of 10 circles represents one recording.

Supplementary Figure S3 shows an example of the temporal changes in the calls’ properties of two brood mates before and during parental visits. Because the calls around parental visits often overlapped, only a small number of calls could be reliably measured and assigned to different individuals. Therefore, it was not possible to assess the calls’ repeatability during parental visits.

## DISCUSSION

The main call of young Scops Owls was found to be more complex than previously described. Koenig (1973) published a few spectrograms but did not report statistics of inter-individual variability. My study has revealed not only fine-scale differences between the individuals’ call spectrograms that were consistent both within and between recordings but also substantial variation in several temporal and spectral measures. The duration of the call and the peak frequency in the ending part of the call had the highest potential for distinctiveness. However, the vocalizations of young Scops Owls exhibited a moderate degree of individuality, which could be insufficient to allow accurate individual recognition. So, what is the origin of this variability, and what is its function?

### A comparison with other species

In nestlings of bird species, various acoustic parameters of food-begging vocalizations exhibit individual variation: pitch (Nakagawa et al. (2001)), call duration (Levréro et al. (2009), Reers and Jacot (2011), Marques et al (2011)), frequency measures among which fundamental frequency and frequency modulation (Levréro et al. (2009), Draganoiu et al. (2006), Kidawa et al. (2023)), spectral entropy (Reers and Jacot (2011), Kidawa et al. (2023)); and proportion of time with harmonic structure (Marques et al (2011)). While the frequency and its modulation are dominant in the literature as identity cues, most authors suggest that individuality is not based on one single component, but on a wide range of components acting simultaneously or in sequence, which presumably reduces discrimination error and reaction time (Rowe 1999).

In my study, two acoustic features stood out for their individual variability and temporal stability: the duration of the call and the acoustic frequency in the ending part of the call. This shows that not only the entire vocalizations but also sub-parts of the vocalizations contain signals that could potentially be used to discriminate between individuals.

While vocal signatures have been found in adults of several species of Strigiformes (Madhavan & Linhart 2024), the only study of individual signatures in young owls is that by Dreiss et al. (2014) of nestling Barn Owls *Tyto alba*. Repeatability estimates of acoustic features found in Barn Owls were comparable with those reported in this study. For example, Call duration was highly repeatable in both species (Scops Owls, average ± SE: 0.82 ± 0.27; Barn Owls, 0.71 ± 0.05). Like in Barn Owl nestlings, Call duration had the highest repeatability than any frequency measure. In contrast, nestling Barn Owls also differed in call rate, but that was not the case in Scops Owls. One important difference is that in Barn Owls vocal distinctiveness was found in the nestling stage. It has been suggested that nestlings recognize each other, and attend to changes in the other’s calling behavior. This could play a role in the context of competition for the food that the parents deliver to them (Dreiss et al. 2014). However, it is not known whether individual signatures are maintained in adult Barn Owls. The Scops Owl is therefore the first owl species that shows some degree of individuality in the young (this study) as well as in the adults (Koenig 1973, Galotti and Sacchi 2001, Dragonetti 2007).

The signature information capacity estimated in this study (H_S_ = 4.48) is comparable to that found in non-colonial and loosely colonial breeding species, such as the Barn Swallow (H_S_ = 4.57; Medvin et al. 1993) and the Tree swallow (H_S_ = 3.20; Leonard et al., 1997). In contrast, higher values of H_S_ have been found in strictly colonial species, for example, the Bank Swallow (H_S_ = 10.2; Beecher, 1991) and the Cliff Swallow (H_S_ = 8.74; Medvin et al. 1993). The statistic H_S_ estimated in this study implies that a maximum of 26 individuals could be potentially identified at the given degree of accuracy. This number corresponds to 6-8 broods of average size present in a certain area, which is not too far from reality in a semi-colonial context like that of this study. Scops Owls are known to form loose colonies with distances between nests as short as 12 m and partially overlapping territories of males (Cramp 1985, Vrezec 2001, Grieco 2018, 2022). Although fledged broods tend not to mix with others, they have been observed entering neighbouring territories (author, unpubl. data). This suggests that the parents face a higher chance of coming in contact with unrelated young, making acoustic signatures and recognition mechanisms more important for this species. Thus, even in territorial birds some degree of intermingling of young after fledging could explain the occurrence of acoustic variability in the offspring (Barg and Mumme, 1994).

A study of the Stone curlew (*Burhinus oedicnemus*; Dragonetti et al. 2013) chick vocalizations shows some similarities with the present study. The Stone Curlew is a non-colonial species that sometimes forms local aggregations with inter-nest distances being below 100 m. When the chicks leave the natal nest and become mobile, the parents face the problem of recognizing their offspring. Dragonetti *et al*. (2013) found that between individual variability of chick calls was higher than that within individuals. However, it only allowed the correct identification of 37% of the subjects (based on LDA; n = 19).

Playback experiments showed that the parent birds responded in a similar way to the calls of their own chicks and that of stranger chicks. The authors concluded that the degree of individuality of the chicks’ voice was too low to enable chick recognition.

### Acoustic distinctiveness in juveniles vs adults

The advertising call (“hoot”) of adult Scops Owls shows remarkable individual variation and stability through time (see the Introduction). Linhart et al. (2022) reported an H_S_ of 7.0 for adult Scops Owls obtained by converting the discrimination score obtained by Galeotti & Sacchi (2001). This indicates that up to 128 adults could potentially be identified using their calls’ distinctive features. So how does the distinctiveness of vocalizations early in life relate to that found in adults? According to Koenig (1973), the hoot of adult Scops Owls derives from the begging calls of the juveniles through a gradual transition, similar to what was found in other species (Redondo & Exposito 1990, Brittan-Powell et al. 1997, Marques et al. 2011). However, Konig (1973) did not provide trajectories of the development of call features. In my study, two acoustic features exhibited marked individual variation: the duration of the entire call and the peak frequency in the ending part of the call. The duration of the call also varies between adults (Galeotti & Sacchi 2001). Therefore, the variability of call duration in the juveniles could reflect the individual trajectories of call duration during development. The peak frequency in the ending part of the call also showed marked individuality and stability. This part of the call could gradually transition to the “plateau” that one can observe in the spectrogram in the adult call (Galeotti & Sacchi 2001, Denac & Trilar 2006, Dragonetti 2007, Grieco 2018). In contrast, the first part of the call of the juveniles has a wide frequency band, and its peak amplitude can occur at different frequencies, also within individuals. This “uncertainty” in this part of the call could reflect the development of the fast frequency downshift at the beginning of the adult call, where the peak frequency drops by several hundred Hz. In short, some features of the begging calls suggest that they are a precursor of the individual variability in the adult.

A contradictory finding was that the time between subsequent calls in individual juveniles was everything but repeatable, both within and between recordings (this study). In contrast, adult Scops Owls are well known for their individual-specific, clockwork-like calling rhythm. In the juvenile, the calling rhythm could be associated with the individual’s current condition, possibly its hunger state, as suggested by the patterns observed during the parental visits. Later in life, this link is presumably lost, and the calling rhythm becomes fixed.

### Possible functions of vocal distinctiveness in juvenile Scops Owls

Individual signatures in adult Scops Owls are probably adaptive because they help mate recognition (Galeotti et al. 1997) or mediate territorial interactions, for example in the recognition of and response to territorial neighbours (Grieco 2022). What could be the net advantage of distinctive characters in food-begging vocalizations early in life?

In the first scenario, the parent could attend to the call distinctiveness to recognize its offspring among a (large) number of juveniles. Offspring recognition is adaptive if the brood is likely to mix with other juveniles of similar age. When the parents return with food, it probably uses a rule of thumb to visit the area where it found the juveniles in the previous visit. Because the juveniles are mobile, the parent still faces the problem of ensuring that it feeds its offspring, not others’. This is particularly the case in breeding aggregations (“loose colonies”), where fledged broods are likely to end up at close distance during their movements. Based on this study, the individuality of the calls of juveniles was moderate.

Both the call classification scores and the information content H_S_ obtained suggest that the parent could potentially recognize its offspring in a group not larger than 20-25 individuals. This figure is probably not far from the number of juveniles that are likely to be found in the area where a fledged brood stays in the first weeks after fledging. For example, in the study site, one can find three to six broods of two to four fledglings in the core area of approximately six hectares. Juveniles move up to 250 m from the natal nest in the first two weeks after fledging (Grieco, in press). It is therefore realistic to assume that, even in the context of semi-colonial breeding, the parents would come in contact with a similar number of other juveniles. The question is whether the degree of individuality found in this study could be sufficient to enable some mechanism of recognition.

If adult Scops Owls can recognize their offspring acoustically, there are other, not necessarily mutually exclusive, functional explanations for the occurrence of acoustic signatures that go beyond the classic parent-offspring recognition problem in a crowded world. The parents could use acoustic signatures to discriminate between their own nestlings and alien (parasitic) nestlings, as found in other species (Colombelli-Négrel et al. 2012, Yang et al. 2015). However, Scops Owls are not subject to any known brood parasites (Mikkola 1983, Cramp 1985), and laying eggs by multiple females in the same nest is a rare event (Bavoux et al. 1991). Therefore, the acoustic distinctiveness of nestlings is unlikely to be of adaptive value in the context of the rejection of unrelated nestlings.

So far vocal distinctiveness has been addressed in the context of parent-offspring communication. In a rather different scenario, young Scops Owls could perceive the identity cues of their siblings, mediating communication between them. This idea is plausible because young Scop Owls spend significant time calling when their parents are absent (Supplementary Figure S3). Vocalizations in the absence of parents are known in other owl species (Roulin et al. 2000, Olsen et al. 2020). Identity cues could help the fledglings to keep contact with each other (Ligout et al. 2016), but they could also play a role in sibling competition over the next feed brought by the parents, as it was found in Barn Owl nestlings (Roulin et al. 2000, Dreiss et al. 2014).

In conclusion, this work has revealed that vocalizations of young Scops Owls show interesting individual variability in their acoustic structure, but it is not clear if that is sufficient to enable individual recognition. This should encourage researchers to address the unanswered questions outlined above and gain insights into the relationships between nesting ecology, social/territorial organization, and communication in this still poorly known species.

## Ethics statement

Fieldwork was performed in accordance with national guidelines and conforms to the European Directive 2010/63/EU.

## Data Availability

Data and code are available at https://figshare.com/s/c2542355eda72bc8faad.

## Supplementary Material

**Supplementary Information S1.pdf**

This document describes the method used to label the calls of young Scops Owls by visual and acoustic inspection, within recordings and between recordings. It also contains Table 1 with a summary of the broods studied and Table 2 with an overview of per-individual classification success in the Linear Discriminant Analysis.

**Supplementary Figure S2.tif**

An example of how the difference in call structure between two individual fledglings spaced apart (A) can be found later when those individuals reunite (B). A: Duration of the calls of two fledglings of the same brood when they were recorded at a distance from each other for about half an hour. The parabolic microphone was pointed to each fledgling alternately and each time the author spoke which side the microphone was pointed to. Because the call given by the animal in focus appeared much louder than that given by the other when plotted as a waveform or spectrogram, it was always possible to establish which of the two individuals made a specific call in the recording. The two vertical lines indicate the moment a parent owl visited the fledglings. B: Duration of the calls of the same brood mates in five subsequent nights, labeled through the visual inspection of the spectrogram (see the patterns indicated in the inset; time scale 1 second). Each circle represents a call in a subset of 10 calls selected for each night. The calls in A were recorded during the last night in B.

**Supplementary Video S3.mp4**

Fragment of a recording of vocalizations of four brood mates (brood 5). The calls were labelled with numbers based on the spectrograms’ visual appearance and acoustic similarity. For details, see Supplementary Information S1. The reader is encouraged to watch this video repeatedly and focus each time on one specific type (e.g. first 1, then 2, etc.).

**Supporting Online Figure S4.tif**

Temporal variation of the calls’ acoustic features in two fledglings of brood 5 during a 31-minute recording that included two visits of adults (indicated by the owl symbols). During this period the owlets were always separated and therefore could be traced. Time is expressed as the time remaining to the first visit (0 s). Each point represents the loudest call selected in a 30-second interval. The thin lines represent the running average calculated with three values per data point. Right: representative spectrograms of the calls at five time points within the recording time (numbers are indicated in the top-left chart). The first spectrogram refers to the individual represented with open circles in the charts.

