## Supplementary Information 1 for "Vocal distinctiveness in young Eurasian Scops Owls (*Otus scops*)"

**Further information about spectrogram classification**

A possible objection to my method is that labelling through the similarity of call spectrograms (and perceived sounds could contain “subjective” elements. Spectrograms that look alike are assigned to the same individual. Because the animals were not marked, the method introduces some chance of error. This issue can be addressed by looking at the calls of single individuals (i.e. when brood size was 1), or siblings that stayed at distance from some time. In both cases there is no chance of identity swaps.

***1. Examples of single fledglings***

Call spectrograms of a single fledgling recorded in the same place over 9 days. This individual was not included in the dataset because of background noise. Note the “~” pattern at about 1.5 kHz.

| July 19, Rec 2885 | July 22, Rec 2929 | July 26, Rec 2973 |
| --- | --- | --- |
| 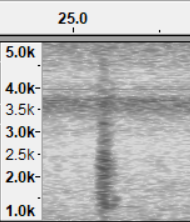 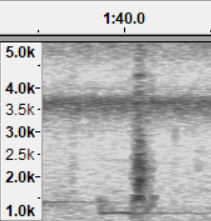 | 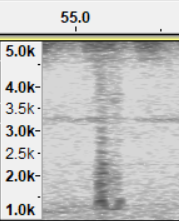 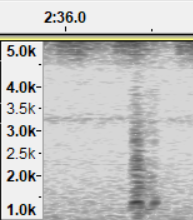 | 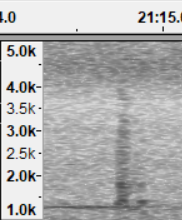 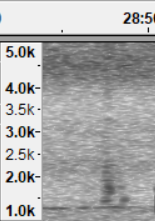 |
| July 26, Rec 2977 (faint)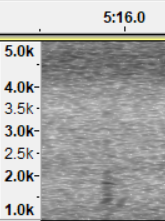 | July 27, Rec 2993  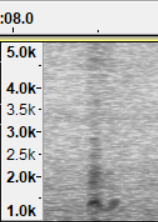 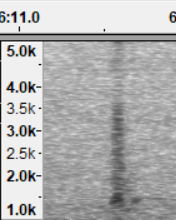 |  |

***2. Calls of fledglings temporarily spaced apart within recordings***

A parabolic microphone amplifies sound that travels directly to the “focus” of the parabolic dish. When pointing the microphone at one bird, the voice of that bird is louder than that of the others. This method was applied for 15 of 23 individuals.

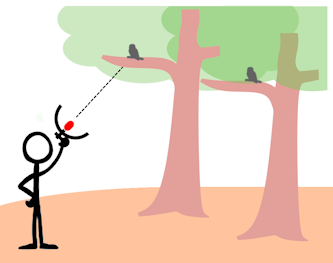

- *Example*. July 24, 2022, brood 6: two young 10 m far apart. One individual was in focus (form “2”).

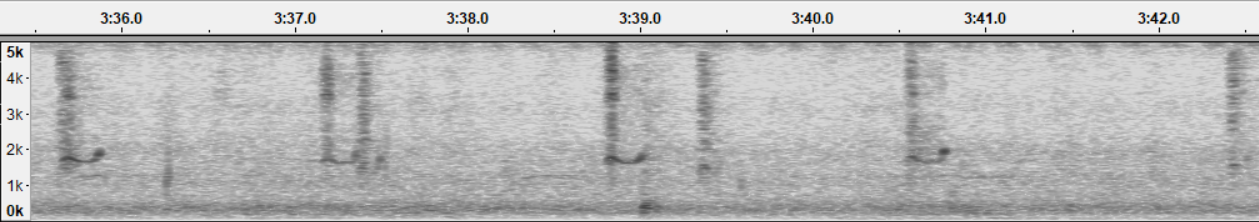

Then, another individual (form “1”, shorter) was in focus, while the other was in the background.

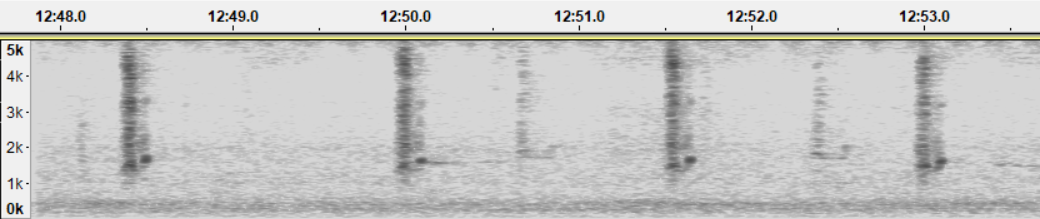

Two days later, form “2” was in focus, while the other was in the background.

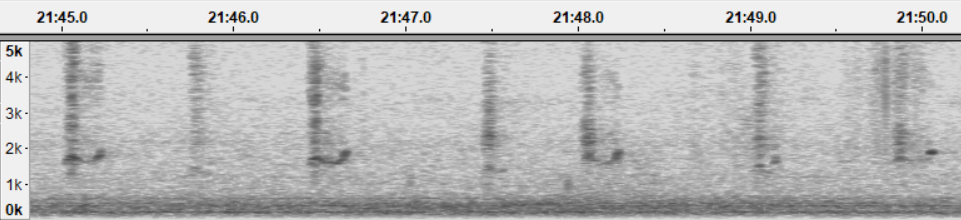

***3. Calls of fledglings temporarily spaced apart over multiple nights***

This is a crucial proof that spectrogram forms are stable over multiple nights and is described in the Results and in Figure 1 of the manuscript.

***4. Examples of stability of forms across recordings***

- *Example 1.* Brood 6, two siblings. Form “1” (top) vs “2” (bottom):

|  | July 24, 2022 | July 25 | July 26 | July 27 | July 28 | July 29 |
| --- | --- | --- | --- | --- | --- | --- |
| **1** | 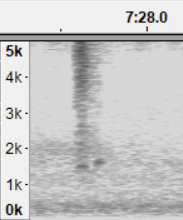 | 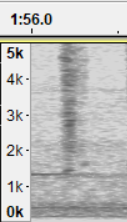 | 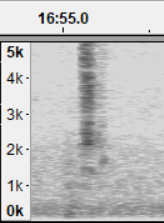 | 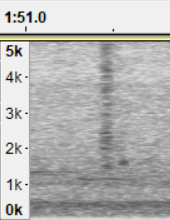 | 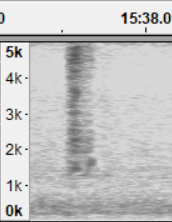 | 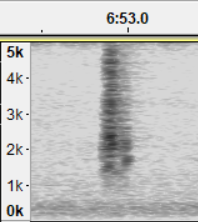 |
| **2** | 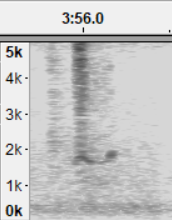 | 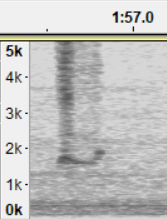 | 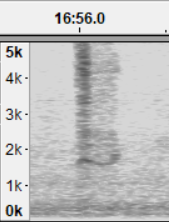 | 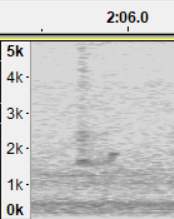 | 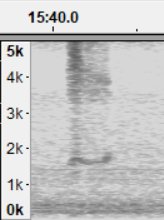 | 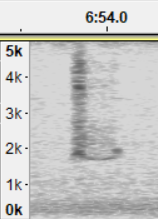 |

- *Example 2*. Brood 5, four owlets. Below: form “1” (with a very clear “S” pattern) vs form “2”. Form “2” was shorter and sounded lower than “1”.

|  | July 25, 2022 | July 26 | July 27 | July 28 | July 29 |
| --- | --- | --- | --- | --- | --- |
| **1** | 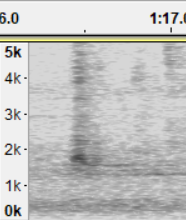 | 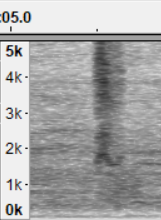 | 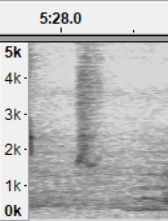 | 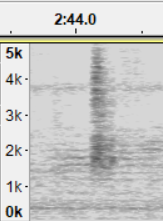 | 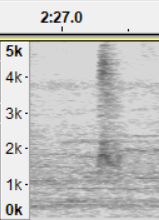 |
| **2** | 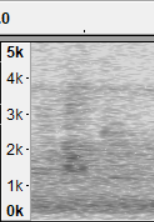 | 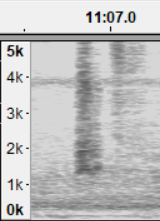 | 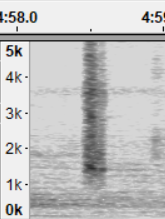 | 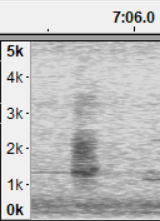 | 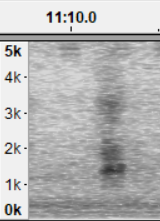 |

See also Supplementary Video S2 for a comparison between the four forms.

***5. Conclusions***

- Single fledglings and fledglings separated for some time produced calls of a consistent form within recordings and across recordings.
- Switching between call forms by the same individual followed continuously was never observed.
- There were always as many forms as the number of fledglings observed.

When integrating the insights exposed above, logic would conclude that an individual owl that was given say label “1” in one recording was the same individual that was given the label “1” in a different recording.

**Call features**

| Call duration (s) | The time from the onset to the end of the call. It was measured from the waveform using the Praat software. |
| --- | --- |
| Time from the previous call (s) | The interval between the onset time of the previous call and that of the current call. It was measured from the waveform using Praat. (1) The start of the call was taken as the start of the first segment of 0.01 s in which the mean Intensity measured in Praat was at least 5 dB greater than in the previous 0.05 s segment. The end of the call was the time at which the local peak in Intensity occurred, or the average time in the case of the two highest local peaks. Sound later than that time was ignored as it could have been reverberation, not the genuine vocalization. |
| PF1/3 (Hz) | The frequency at maximum amplitude in the first third of the call. Peak frequency was measured in Audacity with the function Plot Spectrum for the selected interval. Settings: algorithm Spectrum, function Hanning window, window size 2048. |
| PF3/3 (Hz) | The frequency at maximum amplitude in the last third of the call. Peak frequency was measured in Audacity with the function Plot Spectrum for the selected interval. Settings: algorithm Spectrum, function Hanning window, window size 2048.  Peak frequency 1/3 and Peak Frequency 3/3 were chosen because the initial and the ending parts of the calls were often very different - whereas the initial part was generally noisy, the ending part was more structured. |
| Centre of gravity (Hz) | The average frequency over the entire frequency domain, weighted by the sound amplitude. It was measured with Praat by selecting the call waveform and calling the function Spectrum > View spectral slice, then Query > Get centre of gravity, with power setting 2. |
| Pitch (Hz). | The average pitch across the call duration. It was measured with Praat by selecting the waveform and calling the function Pitch > Get Pitch, with the following settings: frequency range 1000-10000, Analysis method Autocorrelation, Silence threshold 0.03, Voicing threshold 0.1. The intended use of Pitch was to describe the perceived tone of a sound. |
| Average entropy | The average Wiener entropy calculated across the call duration. It was measured with Sound Analysis Pro. Entropy measures the extent to which the sound contains a mixture of frequencies, as opposed to a single frequency or tone. The noisier a sound is, the more energy is smeared within a wide range. Entropy is a pure number and varies between 0 to 1, but here it is reported on a logarithmic scale, ranging between minus infinite (pure tones, complete order) and zero (white noise, complete disorder). |
| Minimum entropy | The minimum Wiener entropy within the call duration (see also Average Entropy above). Minimum Entropy describes the more structured sound, usually found at the end of the call (see an example in Supporting Online Figure S2). |
| Spectrum skewness | A measure of the symmetry of the spectrum. It was measured with Praat by selecting the waveform and calling the function Spectrum > View spectral slice, then Query > Get skewness, with power setting 2. Skewness is a pure number; it is negative when frequencies are concentrated on the right of the spectrum, and positive when the frequencies are concentrated on the left of the spectrum. If skewness is zero, the spectrum is perfectly symmetrical. |

**Supplementary Table S1.**

Basic statistics of the broods analyzed in this study. (*) N: at least one young in the nest. F: all young fledged. (**) Only when fledging was observed. The time from hatching to fledging was estimated to be between 21 and 30 days based on Koenig (1973). Distance indicates the range of the estimated distance between the microphone and birds, valid for all recordings for that brood.

| Brood | Rec. nr. | Date | Nr. of young | Type (*) | Estimated days from hatching (**) | Recording time (min: s) | Calls analyzed | Distance (m) |
| --- | --- | --- | --- | --- | --- | --- | --- | --- |
| 1 |  |  |  |  |  |  |  |  |
|  | 1 | 24-07-2021 | 3 | N | 19-28 | 10:39 | 30 | 4-10 |
| 2 |  |  |  |  |  |  |  |  |
|  | 1 | 06-08-2021 | 3 | F | 31-40 | 07:16 | 30 | 5-10 |
| 3 |  |  |  |  |  |  |  |  |
|  | 1 | 12-07-2022 | 3 | N | 20-29 | 03:48 | 30 | 4-8 |
| 4 |  |  |  |  |  |  |  |  |
|  | 1 | 26-07-2022 | 4 | F | ? | 04:00 | 40 | 15 |
|  | 2 | 29-07-2022 | 4 | F | ? | 03:06 | 40 |  |
| 5 |  |  |  |  |  |  |  |  |
|  | 1 | 25-07-2022 | 4 | F | ? | 03:15 | 40 | 5-10 |
|  | 2 | 26-07-2022 | 4 | F | ? | 13:32 | 40 |  |
|  | 3 | 27-07-2022 | 4 | F | ? | 06:15 | 40 |  |
|  | 4 | 28-07-2022 | 4 | F | ? | 30:04 | 40 |  |
|  | 5 | 29-07-2022 | 4 | F | ? | 13:52 | 40 |  |
| 6 |  |  |  |  |  |  |  |  |
|  | 1 | 20-07-2022 | 3 | N | 19-28 | 33:10 | 30 | 5-12 |
|  | 2 | 22-07-2022 | 3 | F | 21-30 | 31:01 | 30 |  |
|  | 3 | 23-07-2022 | 3 | F | 22-31 | 17:42 | 30 |  |
|  | 4 | 23-07-2022 | 3 | F | 22-31 | 44:13 | 30 |  |
|  | 5 | 24-07-2022 | 2 | F | 23-32 | 40:56 | 20 |  |
|  | 6 | 25-07-2022 | 2 | F | 24-33 | 17:13 | 20 |  |
|  | 7 | 25-07-2022 | 2 | F | 24-33 | 30:09 | 20 |  |
|  | 8 | 26-07-2022 | 2 | F | 25-34 | 42:41 | 20 |  |
|  | 9 | 27-07-2022 | 2 | F | 26-35 | 36:47 | 20 |  |
|  | 10 | 28-07-2022 | 2 | F | 27-36 | 21:55 | 20 |  |
|  | 11 | 28-07-2022 | 2 | F | 27-36 | 32:10 | 20 |  |
|  | 12 | 29-07-2022 | 2 | F | 28-37 | 15:36 | 20 |  |
|  | 13 | 29-07-2022 | 2 | F | 28-37 | 25:58 | 20 |  |
| 7 | 1 | 13-07-2024 | 2 | F | 21-30 | 18:25 | 20 | 5-10 |
|  | 2 | 14-07-2024 | 2 | F | 22-31 | 50:00 | 20 |  |
|  | 3 | 15-07-2024 | 2 | F | 23-32 | 53:00 | 20 |  |
|  | 4 | 16-07-2024 | 2 | F | 24-33 | 48:15 | 20 |  |
|  | 5 | 17-07-2024 | 2 | F | 25-34 | 65:00 | 20 |  |
|  | 6 | 18-07-2024 | 2 | F | 26-35 | 02:00 | 20 |  |
|  | 7 | 19-07-2024 | 2 | F | 27-36 | 06:43 | 20 |  |
| 8 | 1 | 14-07-2024 | 1 | F | 22-31 | 03:07 | 10 | 4-12 |
|  | 2 | 14-07-2024 | 1 | F | 22-31 | 07:34 | 10 |  |
|  | 3 | 15-07-2024 | 1 | F | 23-32 | 11:12 | 10 |  |
|  | 4 | 15-07-2024 | 1 | F | 23-32 | 09:45 | 10 |  |
|  | 5 | 16-07-2024 | 1 | F | 24-33 | 02:25 | 10 |  |
|  | 6 | 16-07-2024 | 1 | F | 24-33 | 08:41 | 10 |  |
|  | 7 | 17-07-2024 | 1 | F | 25-34 | 14:16 | 10 |  |
|  | 8 | 17-07-2024 | 1 | F | 25-34 | 02:37 | 10 |  |
|  | 9 | 18-07-2024 | 1 | F | 26-35 | 05:33 | 10 |  |
|  | 10 | 18-07-2024 | 1 | F | 26-35 | 38:00 | 10 |  |
|  | 11 | 19-07-2024 | 1 | F | 27-36 | 09:27 | 10 |  |
|  | 12 | 19-07-2024 | 1 | F | 27-36 | 33:00 | 10 |  |
|  | 13 | 20-07-2024 | 1 | F | 28-37 | 16:52 | 10 |  |
|  | 14 | 20-07-2024 | 1 | F | 28-37 | 11:46 | 10 |  |
|  | 15 | 21-07-2024 | 1 | F | 29-38 | 07:23 | 10 |  |
|  | 16 | 21-07-2024 | 1 | F | 29-38 | 01:20 | 10 |  |
|  | 17 | 22-07-2024 | 1 | F | 30-39 | 21:00 | 10 |  |
|  | 18 | 22-07-2024 | 1 | F | 30-39 | 56:00 | 10 |  |

**Supplementary Table S2.**

Classification success (%) for individual Scops Owl fledglings based on LDA (see Methods and also Table 1 in this document). The first recording was used for training, and the last recording was used for validation. Each individual was represented with ten calls in both recordings. For example, a classification success of 60 % means that six out of 10 calls of the validation data set were correctly classified for that individual. Top: classification success when the second recording was made one to three days later. Bottom: classification success when the second recording was made six to nine days later. Chance-expected classification success is based on the number of individuals in the sample (across broods) or the average brood size (within broods). (*) P< 0.05, right-tail binomial test. (a) One fledgling of Brood 6 was lost and therefore could not be in the 6–9-day data set. (b) Brood 8 included one fledgling, so the within-brood classification success could not be calculated.

| Brood | Classification success across broods (%) | | | |  | Classification success within broods (%) | | | |
| --- | --- | --- | --- | --- | --- | --- | --- | --- | --- |
| **One/three-day difference** | ID-1 | 2 | 3 | 4 |  | ID-1 | 2 | 3 | 4 |
| Brood 4 (3 days) | 30 | 70 | 0 | 50 |  | 50 | 70 | 0 | 60 |
| Brood 5 (2 days) | 100 | 40 | 100 | 90 |  | 100 | 60 | 100 | 90 |
| Brood 6 (1 days) | 70 | 0 | 80 |  |  | 100 | 70 | 80 |  |
| Brood 7 (3 days) | 10 | 60 |  |  |  | 70 | 100 |  |  |
| Brood 8 (3 days) | 80 |  |  |  |  |  |  |  |  |
| Average vs. expected | 55.7 % vs. 7.1 % * | | | |  | 73.1 vs. 30.8 * | | | |
| **Six/nine-day difference** |  |  |  |  |  |  |  |  |  |
| Brood 6 (9 days) | 80 | 80 | (a) |  |  | 100 | 80 | (a) |  |
| Brood 7 (6 days) | 0 | 80 |  |  |  | 40 | 100 |  |  |
| Brood 8 (8 days) | 70 |  |  |  |  | (b) |  |  |  |
| Average vs. expected | 62.0 % vs. 20.0 % * | | | |  | 80.0 vs. 50.0 * | | | |
