## Supplementary figures and images for "Vocal distinctiveness in young Eurasian Scops Owls (*Otus scops*)"

### Supplementary Figure 2

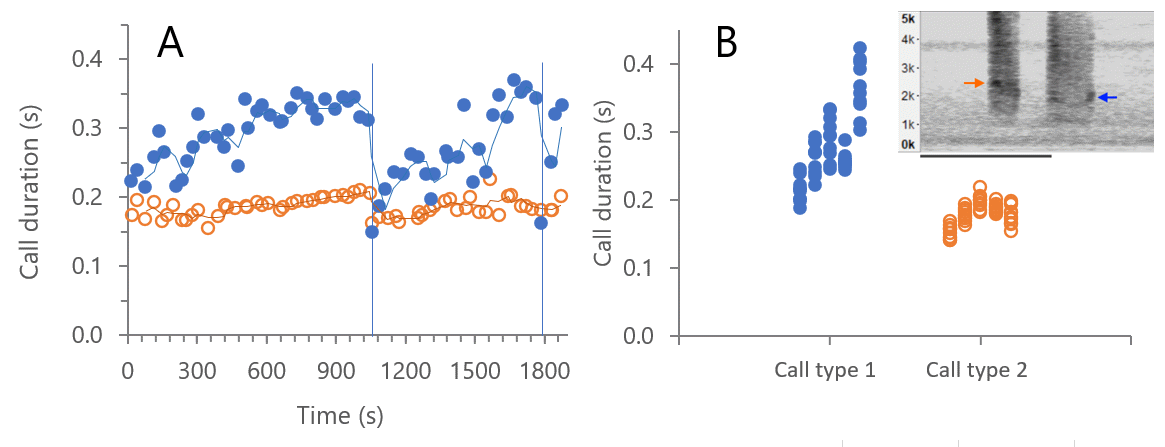

### Supplementary Figure 4

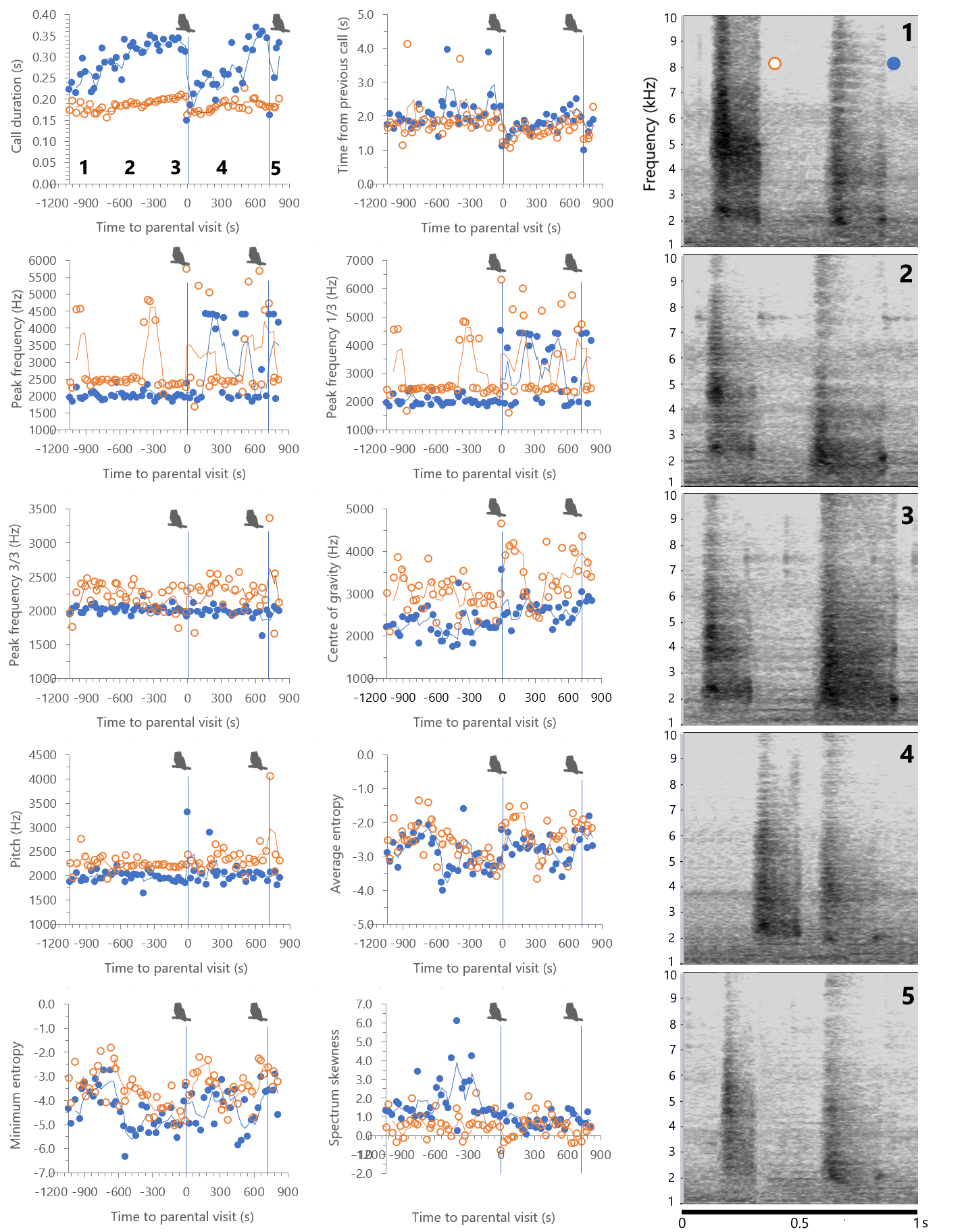
